# Hopeaphenol targets histidine kinase signaling across kingdoms to suppress bacterial virulence and potentiate plant immunity

**DOI:** 10.64898/2026.09.23.753455

**Authors:** Jieun Kang, Nayeon Yoo, Talia Karasov, Venkatesh P Thirumlaikumar, Lydia Story, Eui-Hwan Chung, Aleksandra Skirycz

## Abstract

Small molecules can control plant disease either by disarming the pathogen or by priming host immunity, but single compounds that do both are rare and mechanistically unexplained. Here we show that hopeaphenol (HP), a resveratrol tetramer from stilbene-producing plants, acts on histidine kinase (HK) signaling on both sides of the Arabidopsis– *Pseudomonas* interaction. In *Pseudomonas syringae* pv. *tomato* DC3000, HP represses the type III secretion regulon and motility genes, restricts surface motility independently of effector delivery, and binds a defined subset of virulence-associated sensor kinases while reducing their autophosphorylation; comparable engagement occurs in *Pectobacterium atrosepticum*. Binding and inhibition depend on the tetrameric scaffold rather than the resveratrol monomer. In the host, HP binds the CHASE domains of the cytokinin receptors AHK2, AHK3 and AHK4 and elicits an immune-associated, rather than canonical cytokinin, transcriptional output. HP-dependent potentiation of PTI and ETI responses and early restriction of bacterial growth require AHK3 and AHK4. HP thus coordinates opposing outputs from divergent HK systems across kingdoms.

---

Small molecules can alter both pathogen behavior and host immunity, providing complementary anti-virulence and immune-priming routes to disease control (Conrath et al., 2006; Clatworthy et al., 2007). However, how a single plant-derived compound coordinates these processes remains unclear. Hopeaphenol (HP), a resveratrol tetramer found in stilbene-producing plants, suppresses bacterial virulence without inhibiting growth and enhances Arabidopsis immunity (Kang et al., 2020; Kang et al., 2022a; Kang et al., 2022b). Here we show that HP engages histidine kinase signaling on both sides of Arabidopsis-*Pseudomonas* interaction. This architecture is well characterized in bacterial two-component systems and retained in plants as histidine kinase-based phosphorelays, including Arabidopsis histidine kinase (AHK) cytokinin receptors. In *Pseudomonas*, HP binds virulence-associated sensor kinases and reduces their autophosphorylation. On the host side, HP binds CHASE domains of AHK receptors and selectively facilitates cytokinin signaling associated with plant immunity. These findings establish a cross-kingdom mechanism linking pathogen virulence suppression with host immune potentiation.

Our previous work showed that HP suppresses pathogen virulence while potentiating plant immune responses, but the molecular basis of this dual activity remained unknown (Kang et al., 2020; Kang et al., 2022a; Kang et al., 2022b). To address this, we first defined the bacterial response to HP by transcriptome profiling of *Pseudomonas syringae* pv. *tomato* DC3000 (*Pto* DC3000). HP repressed nearly the entire suite of detected type III secretion system (T3SS)-associated genes, including regulators, secretion-apparatus genes, harpins, chaperones, and *Avr/Hop* effectors (Fig. 1a, Extended Data Fig. 1a). This extends our previous observation of HP-mediated suppression of selected T3SS genes to a regulon-wide response (Kang et al., 2020). In addition, expression of flagellar, chemotaxis, and motility genes was also reduced (Fig. 1b, Extended Data Fig. 1b), and HP strongly restricted surface motility of *Pto* DC3000 (Fig. 1c). This effect remained evident in the effector-deficient ΔCEL and ΔD36E *Pto* DC3000 strains and in the secretion-defective Δ*hrcC* strain, suggesting that reduced motility is not merely a downstream consequence of impaired effector delivery or T3SS apparatus function.

**Fig. 1.**
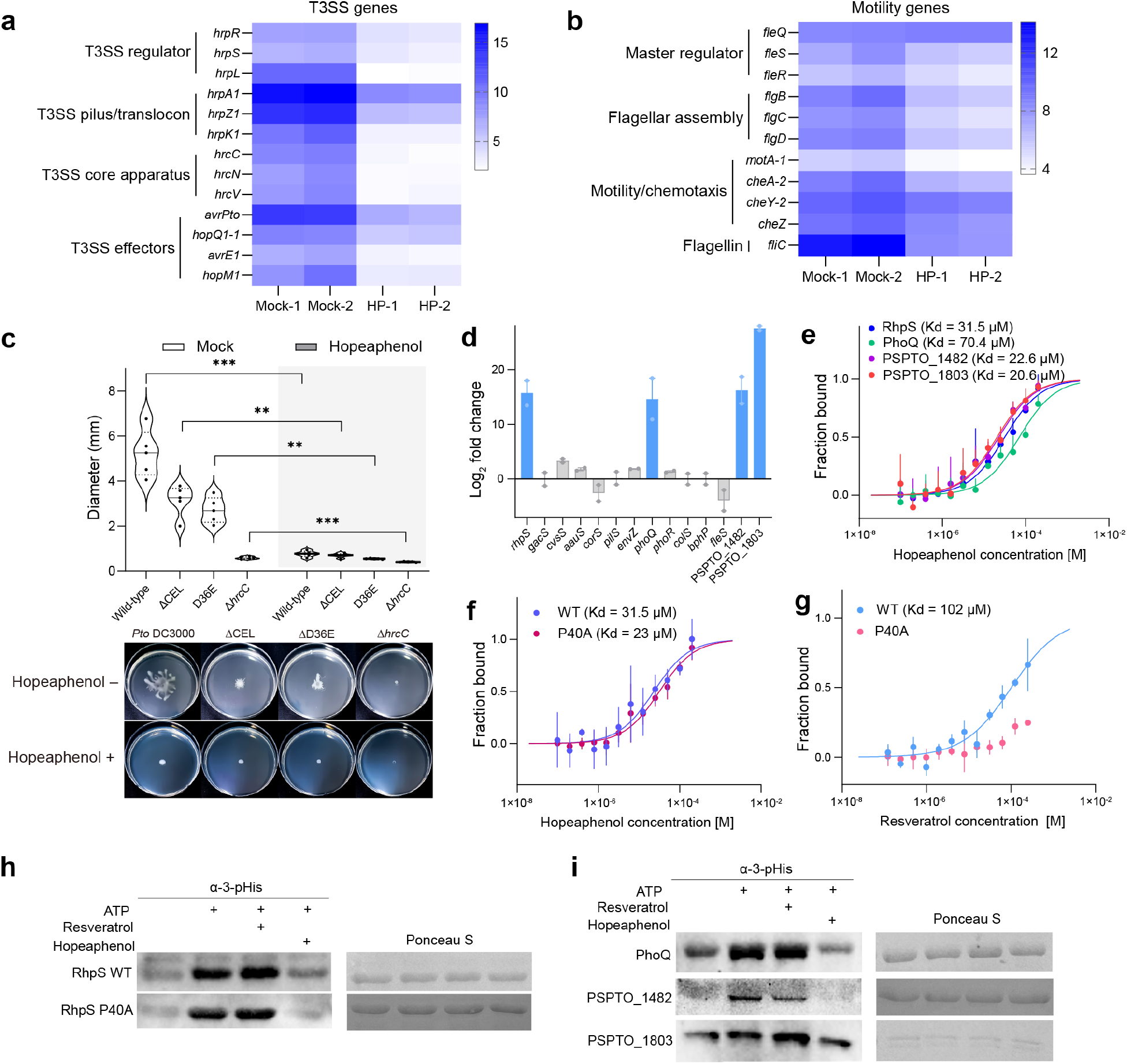
Hopeaphenol represses bacterial virulence by directly targeting *Pto* DC3000 histidine kinases. **a**,**b**, Heat maps of representative T3SS-associated genes (a) and motility-associated genes (b) in *Pto* DC3000 following mock or HP treatment. Genes are grouped by functional categories. Columns represent two independent RNA-seq samples per treatment. Colors indicate log_2_ normalized expression. **c**, Motility phenotype of wild-type *Pto* DC3000 and T3SS-deficient strains ΔCEL, ΔD36E, and Δ*hrcC*. Representative plates are shown below the graph. Violin plots show biological replicates and horizontal lines indicate the median and quartiles (n = 5). Statistical significance was determined by two-sided t-test comparing mock and HP treatment within each strain. \*\**P* < 0.01, \*\*\**P* < 0.001. **d**, Transcript changes of selected *Pto* DC3000 histidine kinase genes following HP treatment relative to mock. *RhpS, PhoQ, PSPTO_1482*, and *PSPTO_1803*, selected for biochemical analysis, are highlighted in blue. Bars represent mean ± SEM from two independent RNA-seq samples. **e**, Microscale thermophoresis (MST) analysis of HP binding to RhpS, PhoQ, PSPTO_1482, and PSPTO_1803. **f**,**g**, MST analysis of HP (f) or resveratrol (g) binding to wild-type RhpS and RhpS P40A mutant. **h, i**, *In vitro* autophosphorylation assays for wild-type RhpS and RhpS P40A mutant (h) and PhoQ, PSPTO_1482, and PSPTO_1803 (i) in the presence of ATP alone or ATP with resveratrol or HP. Phosphohistidine was detected using an anti-3-phosphohistidine antibody. Ponceau S staining shows protein loading control. Blots are representative results from three independent experiments. For MST analyses in **e-g**, points show mean ± SEM from three independent titrations. Solid lines show fitted binding curves.

The coordinated repression of two virulence phenotypes prompted us to examine regulatory pathways that could act upstream of both T3SS and motility. Bacterial two-component systems are major upstream regulators of environmental adaptation and virulence, and typically consist of a sensor histidine kinase and a cognate response regulator that communicate through phosphotransfer (Stock et al., 2000). In *P. syringae*, phosphorylation-dependent regulation of the RhpRS two-component system feeds back on expression of the *rhpRS* operon and is coupled to repression of HrpL-dependent type III secretion system and effector expression (Xie et al., 2020; Huang et al., 2022). RhpRS has also been directly linked to motility in *P. savastanoi* pv. *phaseolicola* 1448A, where phosphorylated RhpR represses *flhA* expression and thereby reduces flagellar biosynthesis and swimming motility (Xie et al., 2019). We therefore reasoned that HP-induced transcriptional changes in histidine kinase genes could identify candidate signaling components associated with HP response. Among HP-responsive *Pto* DC3000 histidine kinases, *RhpS, PhoQ, PSPTO_1482*, and *PSPTO_1803* were among the most strongly induced candidates (Fig. 1d). All four candidate kinases bound HP with comparable micromolar affinities (Kd = 20.6-70.4 μM; Fig. 1e). By contrast, two histidine kinases that were not transcriptionally induced by HP showed no detectable binding (CorS) or substantially weaker binding (GacS, Kd = 275 μM) (Extended Data Fig. 2a). HP likewise did not significantly alter autophosphorylation activity of either CorS or GacS (Extended Data Fig. 2b). Consistent with a degree of target specificity, HP did not bind to or alter the enzymatic activity of pyruvate kinase, a structurally and mechanistically distinct kinase (Extended Data Fig. 2c,d). These results indicate that HP selectively targets a subset of bacterial histidine kinases rather than broadly interacting with the histidine kinase family.

Since HP is a tetrameric resveratrol oligomer and resveratrol oligomerization can generate biological activities and target selectivity distinct from the monomer, we next asked whether histidine kinase recognition reflects the higher-order oligomeric structure of HP rather than a single resveratrol unit (Keylor et al., 2015). A wild type (WT) RhpS bound both resveratrol and HP, enabling a direct comparison of monomer and tetramer recognition (Fig. 1f,g). We therefore tested RhpS P40A variant (P40A), a substitution at a residue previously implicated in polyphenol binding (Xie et al., 2021). P40A strongly weakened resveratrol binding but retained HP binding (Fig. 1f,g). We next asked whether this ligand-specific binding translated into differential effects on kinase activity. HP reduced autophosphorylation of RhpS, RhpS P40A, PhoQ, PSPTO_1482 and PSPTO_1803, whereas resveratrol did not reproduce this broad inhibitory effect under the same conditions (Fig. 1h,i, Extended Data Fig. 3a,b). Together, these results identify multiple bacterial HKs as direct biochemical targets of HP and demonstrate that the tetrameric scaffold confers binding and inhibitory effects distinct from the monomer.

To expand HP engagement of bacterial histidine kinases beyond *Pto* DC3000, we examined *Pectobacterium atrosepticum*, an agriculturally important pathogen that causes potato blackleg and tuber soft rot (Charkowski, 2018). We previously showed that HP suppresses motility, extracellular enzyme production and disease development in *P. atrosepticum* without inhibiting bacterial growth (Kang et al., 2022a). In the corresponding HP-responsive transcriptome, the expression of *ECA_RS4405-ECA_RS4410* response regulator-sensor kinase pair was selectively induced by HP (Extended Data Fig. 3c). ECA_RS4410 bound HP, and the P39A substitution at the position corresponding to RhpS P40 had little effect on HP affinity but markedly reduced resveratrol binding (Extended Data Fig. 3d,e). HP also reduced autophosphorylation of both ECA_RS4410 and ECA_RS4410 P39A (Extended Data Fig. 3f,g). Together, these results show that HP-histidine kinase interactions are not restricted to *Pto* DC3000 and that the binding distinction between HP and resveratrol is retained in a second bacterial phytopathogen, supporting the potential for HP to suppress virulence in both hemi-biotrophic and necrotrophic bacterial pathogens.

Given the effects of HP on bacterial histidine kinases and previously observed enhancement of flg22-triggered reactive oxygen species (ROS) production in Arabidopsis, we next asked whether plant histidine kinases might also contribute to the host response to HP (Kang et al., 2022b). Intriguingly, analysis of the HP-responsive Arabidopsis transcriptome revealed that the cytokinin receptors *AHK2, AHK3*, and *AHK4*, which also belong to the HK family in plant, were significantly induced by HP treatment (Fig. 2a). HP subsequently showed concentration-dependent binding to cyclases/histidine kinases-associated sensory extracellular (CHASE) domains of all three cytokinin receptors (Fig. 2b), while resveratrol did not interact with them under the same conditions (Fig. 2c).

**Fig. 2.**
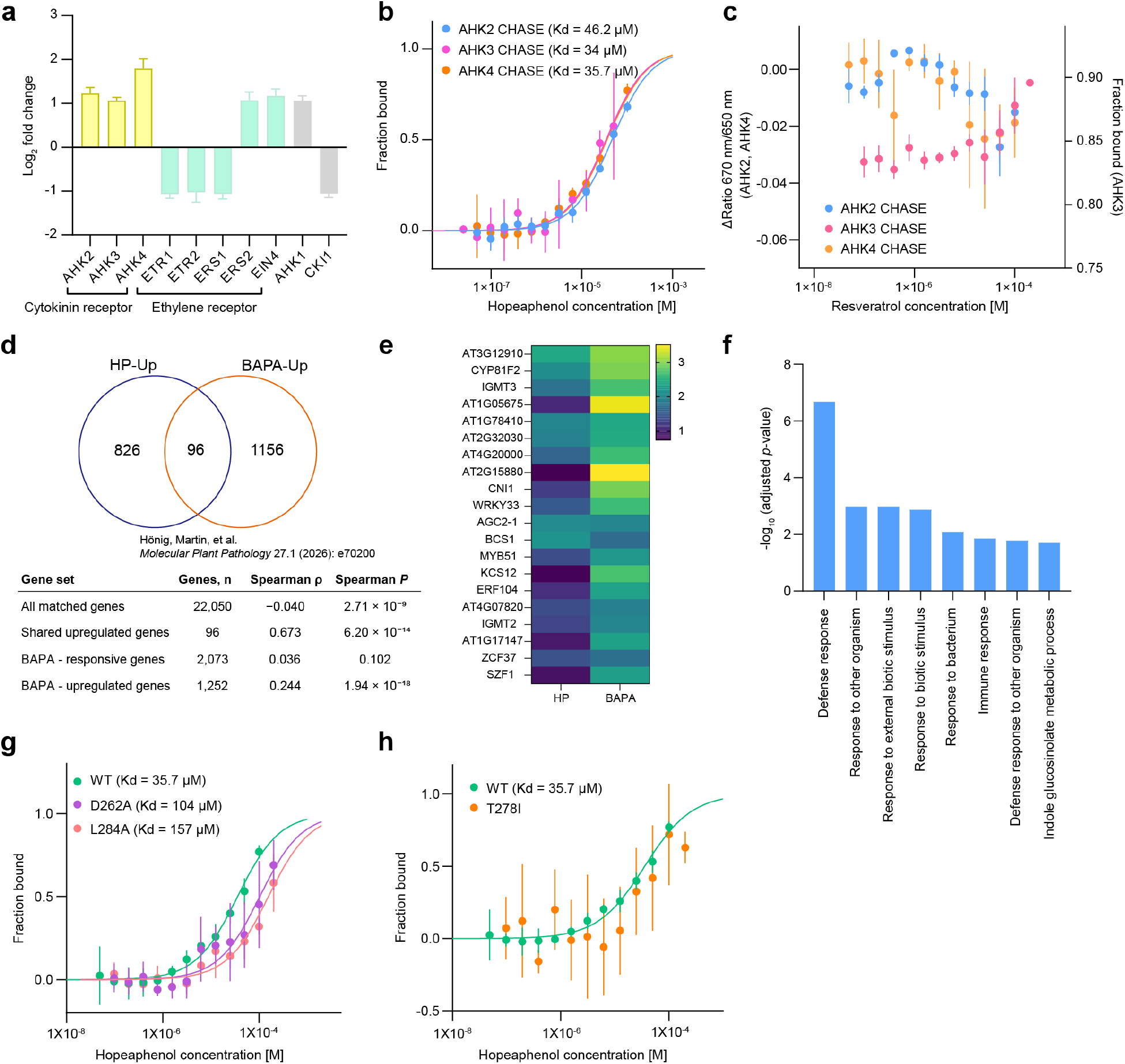
Hopeaphenol interacts with Arabidopsis cytokinin receptors and shares a BAPA-associated immune transcriptional module. **a**, Transcriptional changes of Arabidopsis histidine kinase gene family after HP treatment compared to mock. Cytokinin receptors, ethylene receptors, and other histidine kinase-related genes are indicated. Bars represent mean ± SEM of three independent RNA-seq samples. **b**, MST analysis of HP binding to CHASE domains of Arabidopsis cytokinin receptors AHK2, AHK3, and AHK4. **c**, Binding analysis of resveratrol to CHASE domains of AHK2, AHK3, and AHK4. No reproducible concentration-dependent binding was detected under the conditions tested. **d**, Overlapped genes significantly upregulated by HP and by BAPA. Upregulated genes were defined with adjusted *p* value < 0.05 and log_2_ fold change > 0. The table reports Spearman correlations between HP and BAPA treatments for the indicated gene sets. BAPA dataset was obtained from Hönig et al. (2026). **e**, Heat map of the top 20 genes among 96 genes significantly upregulated by both HP and BAPA, ranked by mean log_2_ fold change across the two treatments. Colors indicate log_2_ fold change relative to the corresponding control. **f**, Gene ontology enrichment analysis of 96 genes significantly upregulated by both HP and BAPA. Bars show −log_10_ (adjusted *p* values) for enriched biological-process terms. **g**, MST analysis of HP binding to wild-type AHK4 CHASE and D262A and L284A variants. **h**, MST analysis of HP binding to wild-type AHK4 CHASE and T278I variant. For MST analyses in **b**,**c**,**g**,**h** points show mean ± SEM from three independent titrations, and solid lines show fitted binding curves.

Cytokinin signaling intersects with immune regulation, but receptor binding alone does not establish canonical cytokinin-like activity (Choi et al., 2010; Argueso et al., 2012; Hönig et al., 2026). We therefore compared the HP-responsive transcriptome with publicly available RNA-seq data from two chemically distinct cytokinin-related compounds with different signaling outputs, tZ (*trans*-zeatin) and the aromatic cytokinin arabinoside BAPA (6-(3-methoxybenzylamino)purine-9-arabinoside) (Hönig et al., 2026). We inferred that BAPA was particularly informative due to its involvement in pathogen resistance dependent on AHK3 while eliciting only limited canonical cytokinin-response gene activation (Hönig et al., 2026). Within the shared upregulated gene sets, HP showed a strong correlation with BAPA than with tZ (Fig. 2d, Extended Data Fig. 4a). 96 genes were significantly upregulated by both HP and BAPA (Spearman ρ = 0.673, Fig. 2d). Among these genes, the top shared genes included defense- and stress-associated regulators, and metabolic genes such as *CYP81F2, IGMT2, IGMT3, WRKY33, MYB51*, and *ERF104* (Fig. 2e). Although these genes were induced by both treatments, BAPA generally produced larger responses among the most strongly induced shared genes (Fig. 2e). Gene ontology (GO) analysis of the shared upregulated gene set highlighted defense response, immune response, response to bacterium, and indole glucosinolate metabolic process (Fig. 2f). However, this similarity was not observed globally, with ρ values of - 0.040 across all matched genes and 0.036 for the broader BAPA-responsive gene set (Fig. 2d). Thus, rather than triggering a broad and generic cytokinin response, HP enhanced the expression of specific immune-related genes typically associated with BAPA. In contrast, HP shared fewer upregulated genes with tZ than with BAPA (49 genes versus 96 genes, Fig. 2d, Extended Data Fig. 4a). Furthermore, the genes shared between HP and tZ were primarily enriched in general stress and stimulus-response categories (Extended Data Fig. 4b,c). We next examined whether HP recognition of AHK4 involves residues known to mediate tZ binding. D262, T278, and L284 are residues of the tZ-binding site of AHK4, and substitution at these positions have previously been shown to impair tZ recognition (Hothorn et al., 2011). In our microscale thermophoresis (MST) assays, AHK4 D262A and L284A variants caused only moderate shifts in HP affinity, whereas the cavity-narrowing T278I variant strongly compromised HP binding (Fig. 2g,h). Together, these transcriptional and binding data indicate that HP engages the cytokinin receptor system differently from tZ, showing immune-associated transcriptional response and a distinct sensitivity to substitutions within the AHK4 tZ-binding site.

We then addressed whether cytokinin receptors are required for HP-induced immunity. HP-dependent ROS enhancement was retained in each single receptor mutant, *ahk2, ahk3*, and *ahk4* (Extended Data Fig. 5a-d). The *ahk2 ahk3* double mutant was excluded from subsequent analysis because its compact growth and markedly smaller leaves under our growth conditions made immune response assays difficult to compare directly with wild-type plants. Among the remaining double mutants, *ahk3 ahk4* clearly lost HP responsiveness and was selected for further plant immunity analyses (Extended Data Fig. 5e-h). In wild-type plants, HP enhanced flg22-triggered ROS accumulation, MAPK phosphorylation and callose deposition (Fig. 3a-c). These HP-dependent increases in ROS, MAPK phosphorylation and callose deposition were lost or strongly attenuated in *ahk3 ahk4* (Fig. 3a-c). HP also potentiated *avrRpt2-* and *avrRpm1-*triggered ion leakage in WT, while this enhancement was reduced in *ahk3 ahk4* (Fig. 3d and Extended Data Fig. 6a). In dexamethasone (Dex)-inducible effector lines, HP pretreatment similarly increased cell death (Extended Data Fig. 6b,c) and enhanced corresponding ROS burst when AvrRpt2 and flg22 responses were co-induced (Fig. 3e). Together, these results support requirement of AHK3 and AHK4 in HP-mediated potentiation of PTI (Pattern-triggered immunity)- and ETI (Effector-triggered immunity)-outputs.

**Fig. 3.**
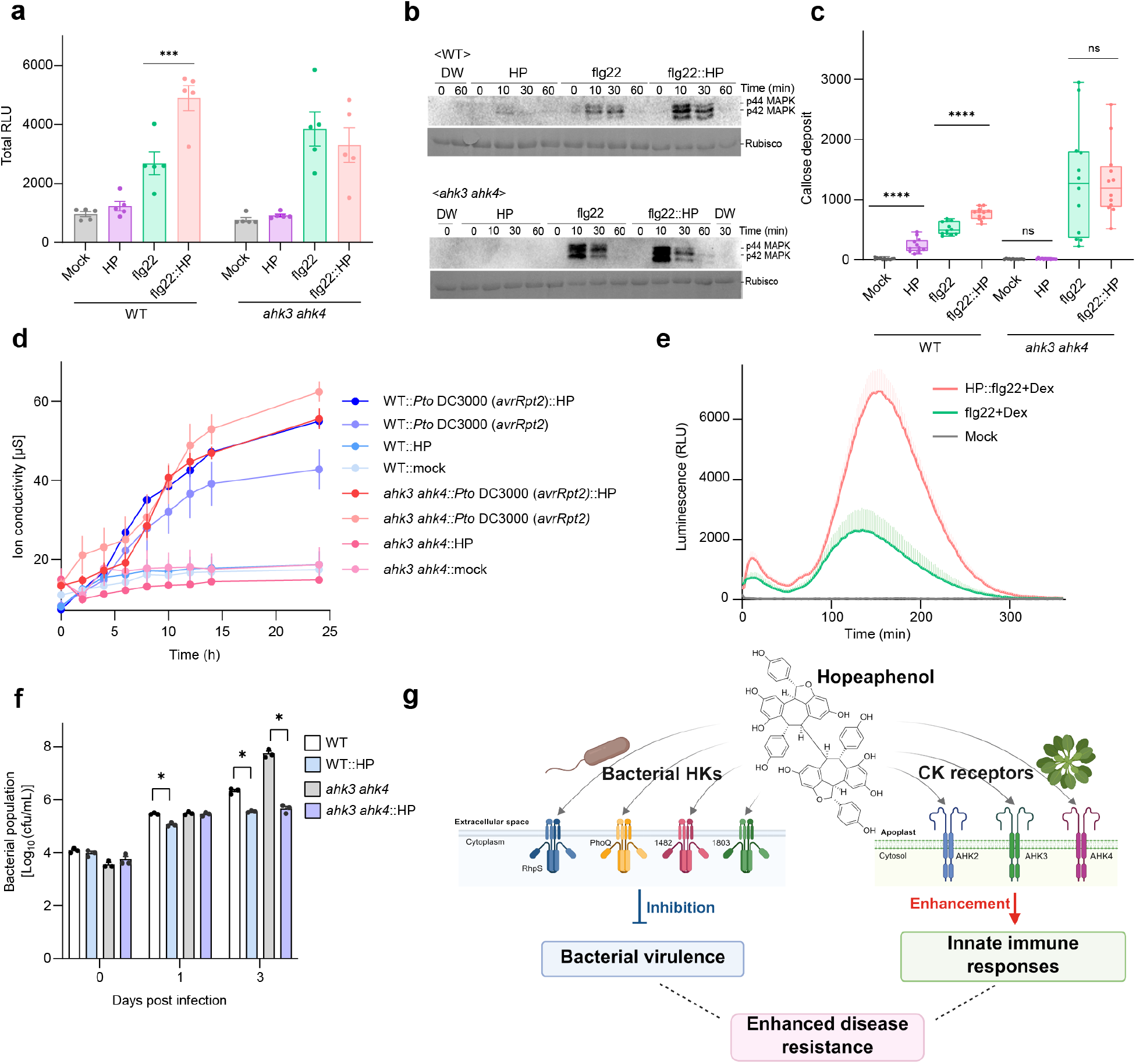
AHK3 and AHK4 are required for hopeaphenol-induced immune potentiation and early bacterial restriction. **a**, Total reactive oxygen species (ROS) production in wild-type and *ahk3 ahk4* leaf discs following mock, HP, flg22, or HP pretreatment followed by flg22. Total ROS is expressed as integrated relative luminescence units (RLU). Bars show mean ± SEM. Dots represent biological replicates (n = 5). Statistical significance was assessed by two-way ANOVA followed by Tukey’s multiple-comparison test. **b**, MAPK activation in WT and *ahk3 ahk4*. Phosphorylated MPK3 and MPK6 were detected by anti-p44/42 antibody. Rubisco staining is shown as loading control. Blots are representative of three independent experiments. **c**, Callose deposition in WT and *ahk3 ahk4*. Boxes show the median and interquartile range, whiskers indicate the minimum and maximum values, and dots represent biological replicates (n = 12). Statistical significance was assessed by two-way ANOVA followed by Tukey’s multiple-comparison test. **d**, Ion leakage in WT and *ahk3 ahk4* following mock, HP, *Pto* DC3000 carrying *avrRpt2*, or HP pretreatment followed by *Pto* DC3000 carrying *avrRpt2*. Points show mean ± SEM (n = 3). **e**, ROS production in Col-0 *Dex::avrRpt2* plants following mock, flg22 plus Dex, or HP pretreatment followed by flg22 plus Dex. Points show mean ± SEM (n = 12). **f**, Growth of *Pto* DC3000 in WT and *ahk3 ahk4* plants pretreated with mock or HP. Bacterial populations were determined at 0, 1 and 3 days post infection. Bars show mean ± SEM. Dots represent biological replicates (n = 3). Statistical significance between mock and HP treatment at each time point was determined using t-test. **g**, Proposed working model of the dual activity of HP. HP directly engages and inhibits bacterial histidine kinase activity, suppressing bacterial virulence, while its association with Arabidopsis cytokinin receptors potentiates immune responses. \**P* < 0.05, \*\*\**P* < 0.001, \*\*\*\**P* < 0.0001, ns; not significant.

The necessity of AHK3 and AHK4 receptors was the most evident during early infection stage. HP reduced the *Pto* DC3000 population at 1 and 3 days post-inoculation in wild-type Col-0 plants, but the early 1-day reduction was compromised in *ahk3 ahk4* double mutant (Fig. 3f). Interestingly, the bacterial growth suppression was nevertheless evident at day 3 in the double mutant (Fig. 3f). These results led us to infer that AHK3/AHK4-dependent host responses contribute particularly to early HP-mediated protection. The recovery of HP-mediated protection at later infection stages in the double mutant may reflect the cumulative impact of direct bacterial virulence suppression (Fig. 1), potentially offsetting the reduced contribution of AHK3/AHK4-dependent host signaling. We therefore propose that HP acts on divergent histidine kinase systems with opposite functional directions in pathogen and host plant: it inhibits a subset of bacterial HKs while engaging cytokinin receptors to potentiate plant defense (Fig. 3g).

Histidine kinase signaling proteins are highly divergent in their signal-sensing domains but share conserved phosphotransfer machinery that transmits these inputs to downstream signaling pathways (Stock et al., 2000; Schaller et al., 2008). Prior strategies for chemically modulating histidine kinases predominantly focused on the conserved catalytic ATP-binding domains to inhibit diverse kinases (Wilke et al., 2015; Velikova et al., 2016; Fihn and Carlson, 2021). In contrast, our study began with HP-induced phenotypes and whole transcriptional responses in both pathogen and host, leading to the elucidation of histidine kinase systems linked to successful plant disease resistance. This phenotype-guided strategy uncovered selective modulation of a restricted set of bacterial histidine kinases and plant cytokinin receptors, rather than general modulation of all histidine kinases. Through its tetrameric resveratrol-derived scaffold, HP acts in two functionally opposite directions: dampening autophosphorylation of selected bacterial histidine kinases to suppress pathogen virulence and engaging cytokinin receptor CHASE domains to potentiate AHK3/AHK4-mediated plant immune responses. This dual activity explains first how a single plant-derived metabolite can weaken pathogen behavior and strengthen host defense simultaneously. More broadly, HP illustrates how a plant-derived oligomer can coordinate divergent signaling systems across kingdoms through a shared histidine kinase signaling architecture rather than obvious sequence or structural conservation. Future work should define the molecular determinants of HP recognition by bacterial histidine kinases and plant cytokinin receptors and determine how these distinct interactions are transmitted into opposing signaling outputs across kingdoms. It will also be important to test whether HP-induced expression of histidine kinases reflects feedback with downstream signaling and whether analogous cross-kingdom regulation extends to other oligomeric polyphenols.

## Methods

### Chemical compound

Hopeaphenol was purified from grape root as described in Kang et al. (2020) and resveratrol was obtained from Sigma-Aldrich. Stock solutions were prepared in acetone and stored at –20°C until use.

### Plant growth

Arabidopsis lines used are listed in Supplementary Table 1. Plants were grown under short-day conditions (9-h light/15-h dark) at 21/18 °C, 60% relative humidity, and a light intensity of 125 μmol m^-2^ s^-1^. Plants aged 4–5 weeks were used for experiments.

### RNA sequencing and transcriptomic analysis

For bacterial transcriptome analysis, single colony of *Pseudomonas syringae* pv. *tomato* DC3000 (*Pto* DC3000) was cultured overnight in King’s B medium. Bacterial culture was transferred to *hrp*-inducing medium Huynh et al. (1989) supplemented with 100 µM hopeaphenol or 0.1% acetone as a control and incubated for 3 h at 18 °C. Cells were harvested, and total RNA was extracted from two independent biological replicates per treatment. RNA libraries were prepared and sequenced on an Illumina platform. Reads were mapped to the *Pto* DC3000 reference genome (GCF_000007805.1_ASM780v1), and gene expression was quantified as RPKM. RPKM values were log_2_-transformed and quantile-normalized, and genes showing at least a twofold change relative to the control were considered HP-responsive. Raw and processed sequencing data, together with full processing details, are available through the Gene Expression Omnibus (GEO) under accession GSE341978.

For plant transcriptome analysis, wild-type *Arabidopsis thaliana* Col-0 plants were pretreated with hopeaphenol or 0.1% acetone for 16 h and subsequently infiltrated with *Pto* DC3000 at 1 × 10^6^ CFU ml^−^1. Infiltrated leaves were collected 3 h after inoculation and immediately frozen in liquid nitrogen. Three independent biological replicates were collected for each condition. Total RNA was extracted, and stranded mRNA libraries were sequenced on Illumina platform. Reads were mapped to Arabidopsis TAIR10.1 reference genome, and differential expression was analyzed using DESeq2. Genes with an adjusted *P* value < 0.05 and an absolute log_2_ fold change ≥ 1 were considered differentially expressed. Raw and processed data are available in GEO under accession GSE341977. HP-responsive genes were compared with published BAPA- and *trans*-zeatin-responsive gene sets by matching AGI identifiers. Same-direction overlaps were determined using an adjusted *P* value < 0.05 and log_2_ fold change > 0, and expression concordance was assessed using Spearman correlation. Gene ontology enrichment analysis was performed for the shared upregulated gene sets using ShinyGO 0.85.

### Motility assay

Overnight cultures of the indicated *Pto* DC3000 strains (Supplementary Table 2) were adjusted to an OD_600_ of 2.0, and 2 µl of each culture was spotted onto swarming agar medium containing 0.5% peptone, 0.3% yeast extract and 0.4% agar. Plates were supplemented with 100 µM hopeaphenol and incubated at 28 °C. Motility was monitored for 48 h, and plates were photographed at 48 h.

### Plasmid construction

Coding sequences of the indicated bacterial histidine kinases and AHK3 CHASE domain (Supplementary Table 3) were amplified and cloned into pENTR/D-TOPO, followed by transfer into pDEST-HisMBP expression vector (Addgene #11085) using Gateway LR recombination (Supplementary Table 4). Sequences coding AHK2 and AHK4 CHASE domains were synthesized in pTwistENTR by Twist Bioscience and transferred into pDEST-HisMBP. All constructs were verified by sequencing.

### Protein purification

Recombinant proteins listed in Supplementary Table 4 were produced in *Escherichia coli* BL21 using terrific broth autoinduction medium. Frozen bacterial pellets were resuspended in equilibration buffer, lysed by sonication and processed for purification. The soluble lysates were loaded onto a 1-ml HisTrap column (Cytiva) connected to an NGC Quest 10 chromatography system (Bio-Rad). The column was equilibrated with 25 mM Tris–HCl, pH 8.0, 300 mM NaCl, 5% (v/v) glycerol and 10 mM imidazole, washed with 10 column volumes of the same buffer, and proteins were eluted using a linear gradient of 10–300 mM imidazole. Fractions containing the target protein were pooled, concentrated and buffer-exchanged into 100 mM Tris–HCl, pH 7.5, using centrifugal filters. Proteins were further purified by size-exclusion chromatography using a Superdex 200 Increase 10/300 GL column (Cytiva) equilibrated with Phosphate-Buffered Saline (PBS) containing 137 mM NaCl, 2.7 mM KCl, 10 mM Na_2_HPO_4_ and 1.8 mM KH_2_PO_4_, pH 7.4, and operated at a flow rate of 0.45 ml min^-1^ for 30 min under isocratic conditions. Fractions containing the target protein were pooled and stored at 4 °C until use.

### Site-directed mutagenesis

Site-directed mutagenesis was performed using the Q5 Site-Directed Mutagenesis Kit (New England Biolabs) according to the manufacturer’s instructions. Mutagenic primers were designed using the NEBaseChanger online tool and are listed in Supplementary Table 3. The pHisMBP_RhpS, pHisMBP_4410 and pHisMBP_CHASE4 constructs were used as templates to generate RhpS P40A, ECA_RS4410 P39A and the AHK4 CHASE-domain variants D262A, T278I and L284A, respectively. All mutant constructs were verified by sequencing, and recombinant proteins were expressed and purified as described above.

### Microscale thermophoresis (MST)

Interactions between recombinant proteins and hopeaphenol or resveratrol were measured by MST using a Monolith instrument (NanoTemper Technologies). His-tagged proteins were fluorescently labelled with RED-tris-NTA dye according to the manufacturer’s instructions. Each compound was prepared as a 16-point twofold dilution series in PBS. Binding curves were generated from normalized fluorescence signals, and dissociation constants were estimated using Monolith Analysis software. Fraction-bound values or Δratio 670 nm/650 nm were used to compare independent titrations. Three independent titrations were performed for each protein–ligand pair.

### Autophosphorylation assay

Purified histidine kinases (2–10 µM) were incubated with 2.5 mM ATP in a buffer containing 50 mM Tris–HCl, pH 7.8, 100 mM NaCl, 10 mM MgCl_2_ and 1 mM DTT for 15 min at 25 °C. Hopeaphenol or resveratrol was included at a final concentration of 100 µM where indicated. Reactions were terminated by adding 1X SDS sample buffer, and histidine phosphorylation was detected by immunoblotting with an anti-N3-phosphohistidine (3-pHis) antibody, clone SC56-2 (Sigma). For dephosphorylation assays, the autophosphorylation reaction was performed in the presence of 13 µM phosphohistidine phosphatase 1 (PHPT1). Protein loading was assessed by Ponceau S staining, and phosphorylation signals were quantified relative to the corresponding protein-loading signal.

### NADH-coupled pyruvate kinase activity assay

NADH-coupled pyruvate kinase activity was adapted from Lee et al. (2023). Each reaction contained 0.5 mM NADH, 2 mM phosphoenolpyruvate, 5 mM ATP, 2 unit pyruvate kinase M1 (PKM1) and 4 unit lactate dehydrogenase in assay buffer containing 50 mM Tris–HCl, pH 7.8, 100 mM NaCl, 25 mM reduced glutathione and 25 mM MgCl_2_. Hopeaphenol or resveratrol was added at a final concentration of 100 µM. NADH oxidation was monitored by measuring absorbance at 340 nm over time. Absorbance values were converted to NADH amounts using a calibration curve, and kinase activity was calculated from the linear rate of NADH consumption and expressed as nmol NADH consumed per second.

### NADH calibration curve

NADH calibration curve was generated by measuring the absorbance at 340 nm of 0, 0.039, 0.078, 0.156, 0.313, 0.625 and 1.25 mM NADH prepared in the buffer containing 50 mM Tris–HCl, pH 7.8, and 100 mM NaCl. The resulting linear regression was used to convert absorbance values from the enzyme-coupled assay into NADH amounts.

### ROS assay

Arabidopsis leaves were infiltrated with 100 μM hopeaphenol or 0.1% acetone in water as a mock control. After 16 h, leaf discs were collected using No. 2 cork borer and floated in 200 μl sterile water in white 96-well plates for 3 h at room temperature under continuous illumination. The water was removed and replaced with 200 μl reaction solution containing 100 nM flg22, 0.2 mM luminol and 20 μg ml^-1^ horseradish peroxidase. For *Dex::avrRpt2* line, dexamethasone was added at 5 μM to induce *AvrRpt2* expression, either alone or together with flg22 as indicated. Luminescence was measured every minute using a VANTAStar microplate reader (BMG LabTech). ROS production was expressed as relative light units over time or as total photon counts integrated over the measurement period.

### MAPK phosphorylation

Arabidopsis leaves were pretreated with 100 μM hopeaphenol or 0.1% acetone mock solution for 16 h and subsequently infiltrated with 100 nM flg22. Samples were collected at 0, 10, 30 and 60 min after flg22 treatment. Four leaf discs were collected for each treatment and homogenized in extraction buffer containing 20 mM Tris-HCl, pH 8.0, 150 mM NaCl, 1 mM EDTA, 1% Triton X-100, 0.1% SDS, 10 mM DTT, 1X protease inhibitor cocktail and 1X phosphatase inhibitor cocktail. Aliquots of 60 μl protein extract were mixed with 20 μl of 6X SDS sample buffer and analyzed by SDS-PAGE and immunoblotting. Activated MAPKs were detected by anti-phospho-p44/42 MAPK (Erk1/2, Thr202/Tyr204) antibody (Cell signaling Technology).

### Callose deposition assay

Arabidopsis leaves were infiltrated with 100 μM hopeaphenol or 0.1% acetone mock solution for 16 h and challenged with 100 nM flg22. After an additional 16 h, leaf discs were collected and cleared in ethanol to remove chlorophyll. Samples were rehydrated in distilled water for 30 min and stained overnight with 0.1% aniline blue prepared in 0.1 M K_2_HPO_4_. Stained leaf discs were mounted in 50% glycerol and imaged using a Nikon C2 confocal microscope. Callose deposits were quantified using ImageJ.

### Ion conductivity

Arabidopsis leaves were infiltrated with *Pto* DC3000 carrying *avrRpt2* or *avrRpm1* at an OD_600_ of 0.1. Hopeaphenol was infiltrated 16 h before bacterial inoculation. Four leaf discs were collected for each treatment, immersed in 6 ml distilled water and incubated under continuous light. Ion conductivity was measured every 2 h using a conductivity meter. For assays using *Dex::avrRpt2* and *Dex::avrRpm1* lines, leaf discs were incubated in 20 μM dexamethasone solution, and conductivity was monitored under the same conditions.

### *In planta* bacterial growth

Arabidopsis plants were infiltrated with 100 μM hopeaphenol or 0.1% acetone solution 16 h before bacterial inoculation. *Pto* DC3000 was resuspended in 10 mM MgCl_2_ at 1 × 10^5^ CFU ml^-1^ and infiltrated into the abaxial side of leaves using needleless syringe. At 0,1 and 3 days post-inoculation, four leaf discs were collected for each treatment and homogenized in 10 mM MgCl_2_. Serial dilutions of the lysates were plated on King’s B agar and incubated at 28 °C until colonies became visible. Bacterial populations were calculated from colony counts and expressed as log_10_ CFU cm^-2^.

## Data availability

The RNA-sequencing data generated in this study are available in GEO under accession numbers GSE341977 and GSE341978. The data that support the findings of this study are available from the corresponding authors upon request.

## Acknowledgements

The authors would like to acknowledge the support from Michigan State University and Korea University. We thank Elliot Meyerowitz for providing Arabidopsis cytokinin receptor double-mutant lines and Tatsuo Kakimoto for originally generating these lines and generously making them available to the research community.

## Author contributions

J.K., N.Y., T.K., E.-H.C. and A.S. designed the experiments. J.K., N.Y., V.P.T. and L.S. performed the experiments. J.K., E.-H.C. and A.S. wrote the manuscript.

