## Extended Data Figures for "Hopeaphenol targets histidine kinase signaling across kingdoms to suppress bacterial virulence and potentiate plant immunity"

**Extended Data Fig. 1**

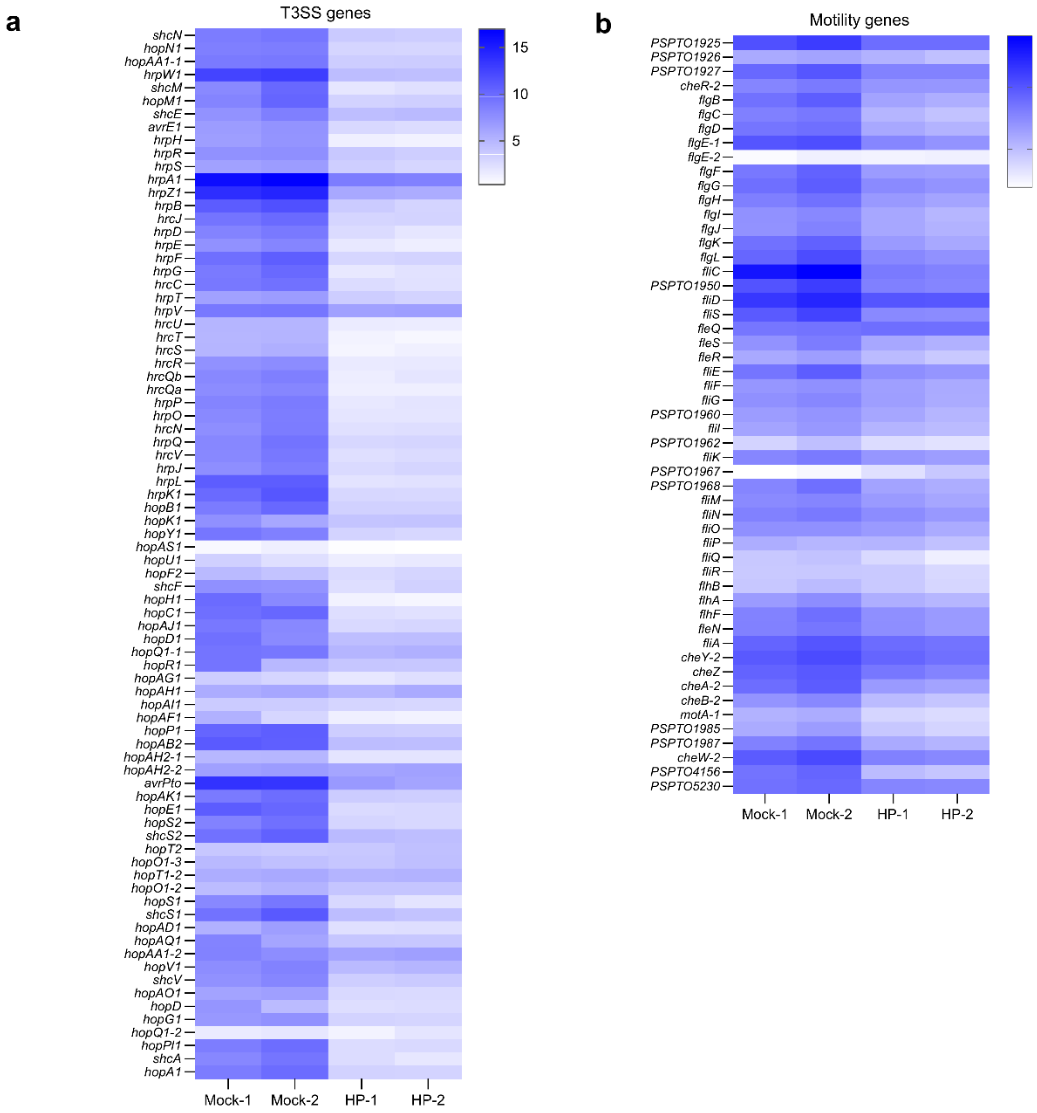


**Extended Data Fig. 1. HP broadly represses T3SS and motility-associated gene expression in *Pto* DC3000. a,b,** Heat maps showing full detected set of T3SS-associated genes (a) and motility-associated genes (b) in *Pto* DC3000. Columns represent two independent RNA-seq samples following mock or HP treatment. Colors indicate $log_2$ normalized expression.

**Extended Data Fig. 2**

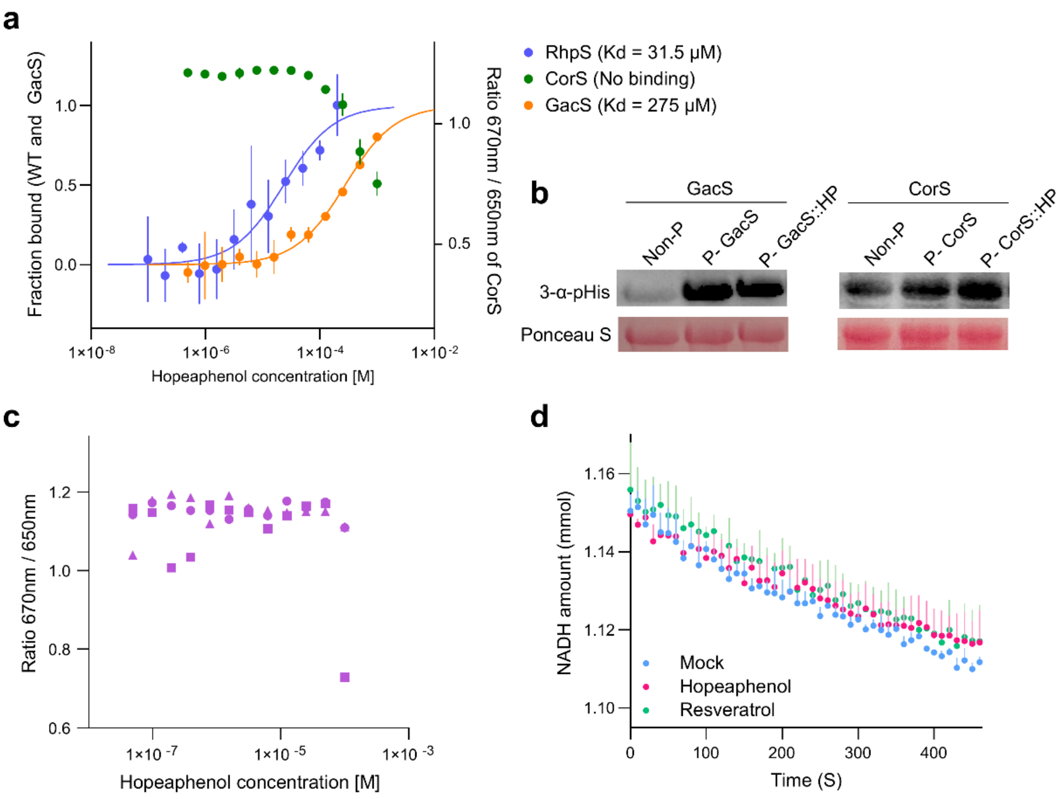


**Extended Data Fig. 2. HP selectively binds and inhibits a subset of bacterial histidine kinases. a,** MST analysis of HP to RhpS and non-HP-induced histidine kinases CorS and GacS. GacS showed substantially weaker binding (Kd = 275 μM) to HP than RhpS (Kd = 31.5 μM), whereas no detectable binding was observed for CorS. Points show mean ± SEM from three independent titrations, and solid lines show fitted binding curves. **b,** *In vitro* autophosphorylation assays of GacS and CorS in the absence or presence of HP. Phosphohistidine was detected using an anti-3-phosphohistidine antibody, and Ponceau S staining shows protein loading. Blots are representative of three independent experiments. **c,** MST analysis of HP binding to pyruvate kinase. No reproducible concentration-dependent binding was detected. Independent titrations are indicated by different symbols. **d,** Pyruvate kinase activity in the presence of mock, HP or resveratrol, measured using an NADH-coupled enzymatic assay. Points show mean ± SEM from two independent experiments.

**Extended Data Fig. 3**

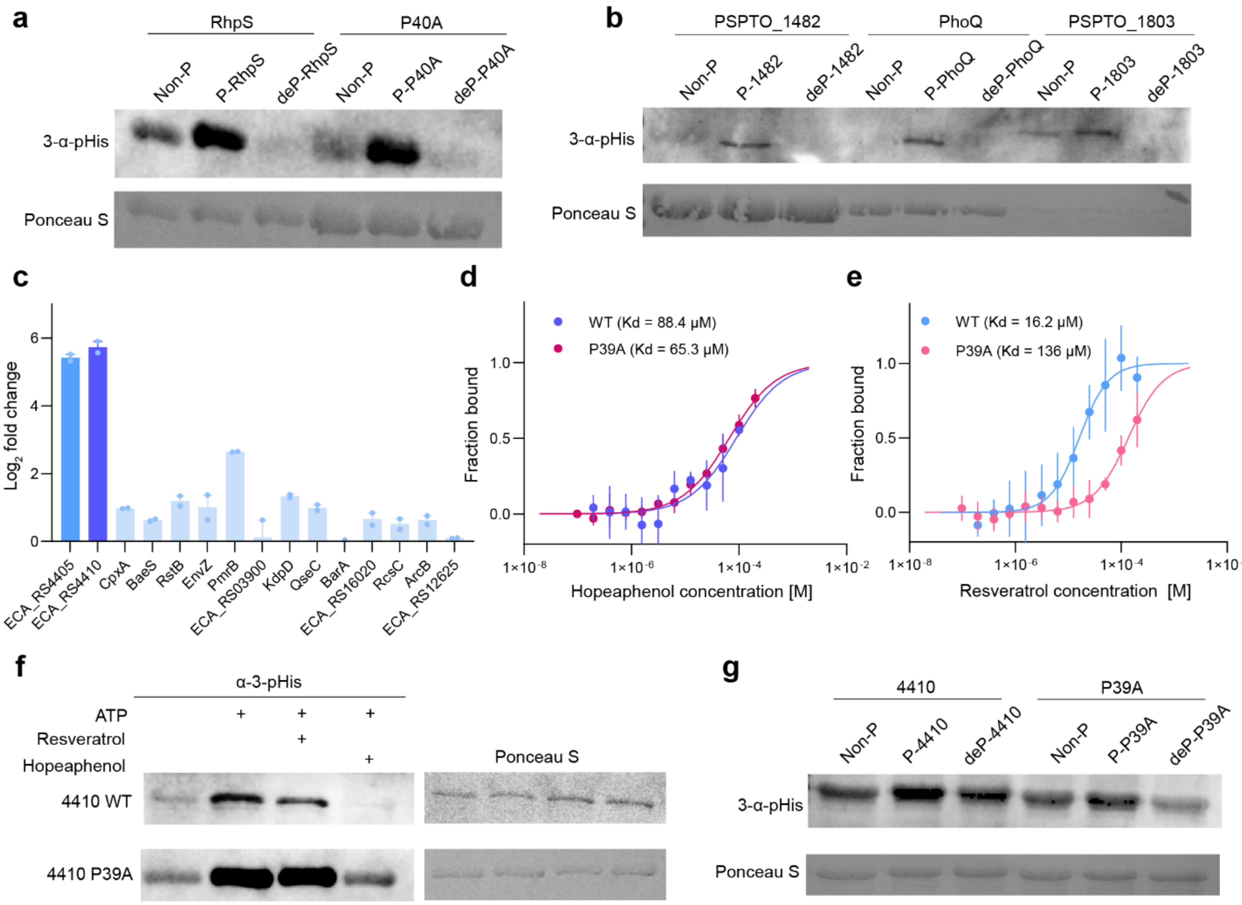


**Extended Data Fig. 3. Phosphohistidine validation and biochemical analysis of HP-responsive *Pto* DC3000 and *P. atrosepticum* histidine kinases. a,b,** Validation of phosphohistidine detection for RhpS and RhpS P40A (a) and PhoQ, PSPTO_1482, and PSPTO_1803 (b). Recombinant proteins were analyzed in their non-phosphorylated (Non-P), ATP-autophosphorylated (P) and dephosphorylated (deP) states using anti-3-phosphohistidine antibody. **c,** Transcript changes of selected two-component system genes in HP-treated *P. atrosepticum* relative to mock. *ECA_RS4405* and *ECA_RS4410* are highlighted. Bars show mean ± SEM from two independent RNA-seq samples. Data are from Kang et al. (2022). **d,e,** MST analysis of HP (d) or resveratrol (e) binding to wild-type ECA_RS4410 and ECA_RS4410 P39A. P39 corresponds to RhpS P40 by sequence alignment. Points show mean ± SEM from three independent titrations, and lines show fitted binding curves. **f,** *In vitro* autophosphorylation assay for wild-type ECA_RS4410 and P39A in the presence of ATP alone or ATP with resveratrol or HP. Phosphohistidine was detected using anti-3-phosphohistidine antibody. **g,** Validation of phosphohistidine detection for wild-type ECA_RS4410 and P39A in their non-phosphorylated, ATP-autophosphorylated and dephosphorylated states. All blots (a,b,f,g) are representative of three independent experiments and Ponceau S staining shows protein loading control.

**Extended Data Fig. 4**

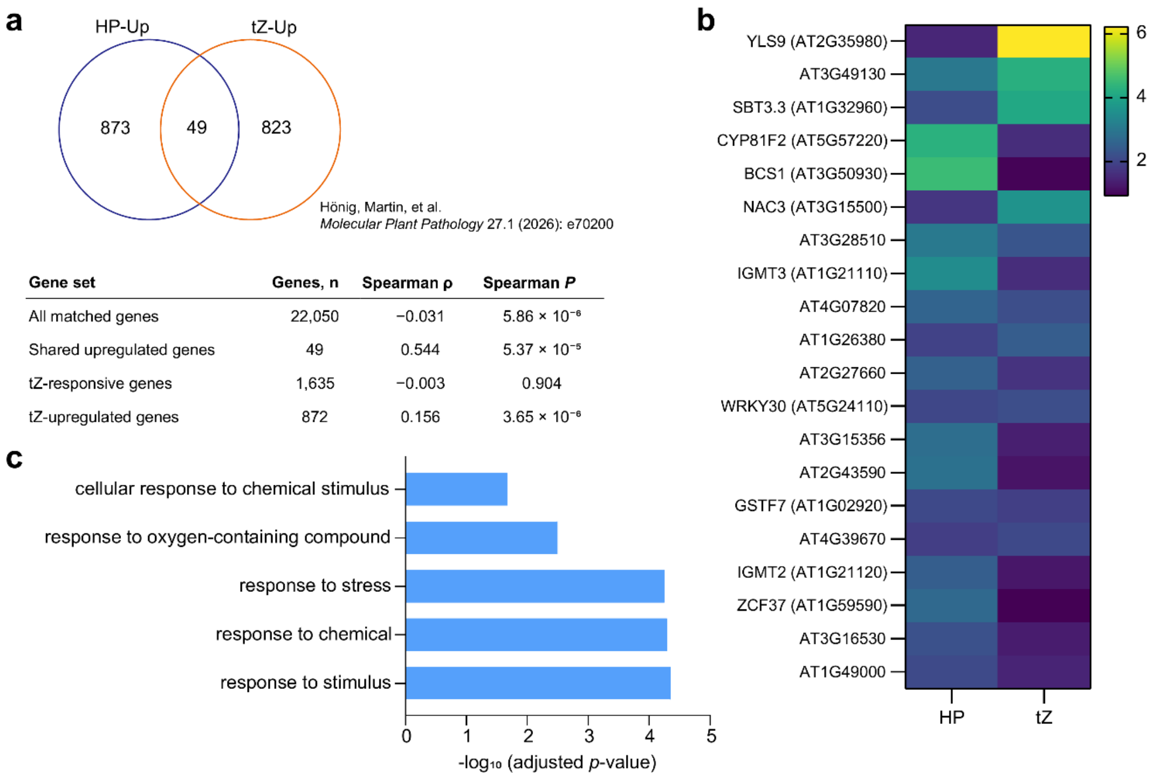


| Gene set | Genes, n | Spearman ρ | Spearman *P* |
|---|---|---|---|
| All matched genes | 22,050 | −0.031 | $5.86 \times 10^{-6}$ |
| Shared upregulated genes | 49 | 0.544 | $5.37 \times 10^{-5}$ |
| tZ-responsive genes | 1,635 | −0.003 | 0.904 |
| tZ-upregulated genes | 872 | 0.156 | $3.65 \times 10^{-6}$ |

**Extended Data Fig. 4. HP shows limited overlap with the canonical *trans*-zeatin transcriptional response. a,** Overlap between genes significantly upregulated by HP and by tZ. Upregulated genes were defined as adjusted *p* value < 0.05 and $\log_2$ fold change >0. The table reports Spearman correlations between HP- and tZ-induced $\log_2$ fold change for the indicated gene sets. tZ dataset was obtained from Hönig et al. (2026). **b,** Heat map of the top 20 genes among 49 genes significantly upregulated by both HP and tZ. Colors indicate $\log_2$ fold change relative to the corresponding control. **c,** Gene ontology enrichment analysis of 49 genes significantly upregulated by both HP and tZ.

**Extended Data Fig. 5**

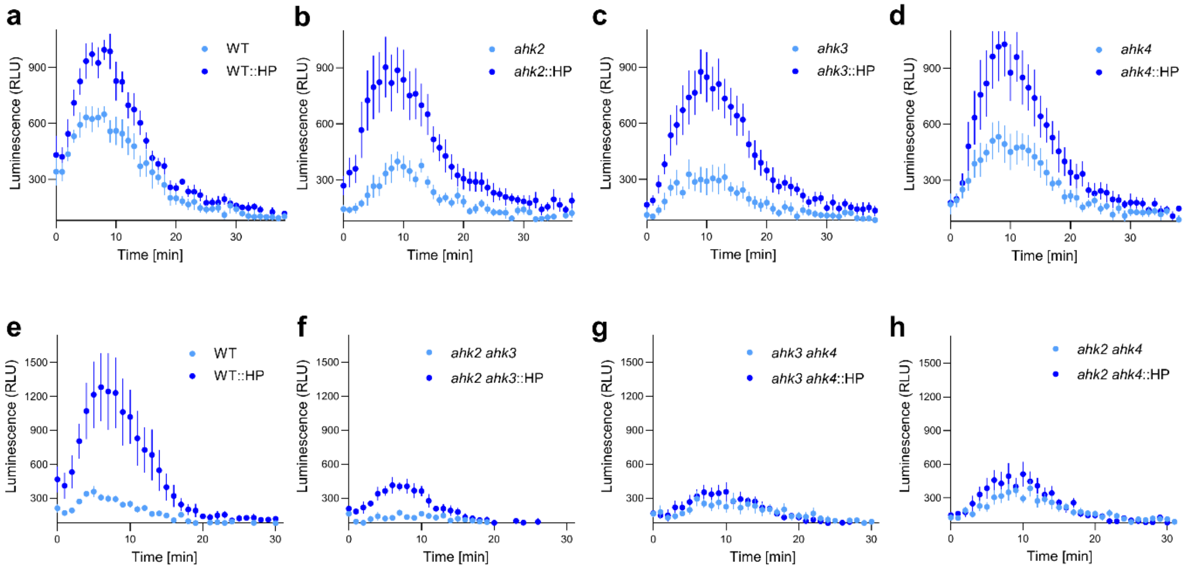


**Extended Data Fig. 5. Cytokinin receptor mutants differ in HP-dependent ROS enhancement. a-d,** flg22-triggered ROS production in wild-type (a) and the single cytokinin receptor mutants *ahk2* (b), *ahk3* (c) and *ahk4* (d) following mock or HP pretreatment. **e-h,** flg22-triggered ROS production in WT (e) and the double cytokinin receptor mutants *ahk2 ahk3* (f), *ahk3 ahk4* (g) and *ahk2 ahk4* (h) following mock or HP pretreatment. Panels a-d were obtained in one experiment, whereas panels e-h were obtained in a separate experiment, each with its corresponding wild-type control (a and e). Points show mean ± SEM, n = 6.

**Extended Data Fig. 6**

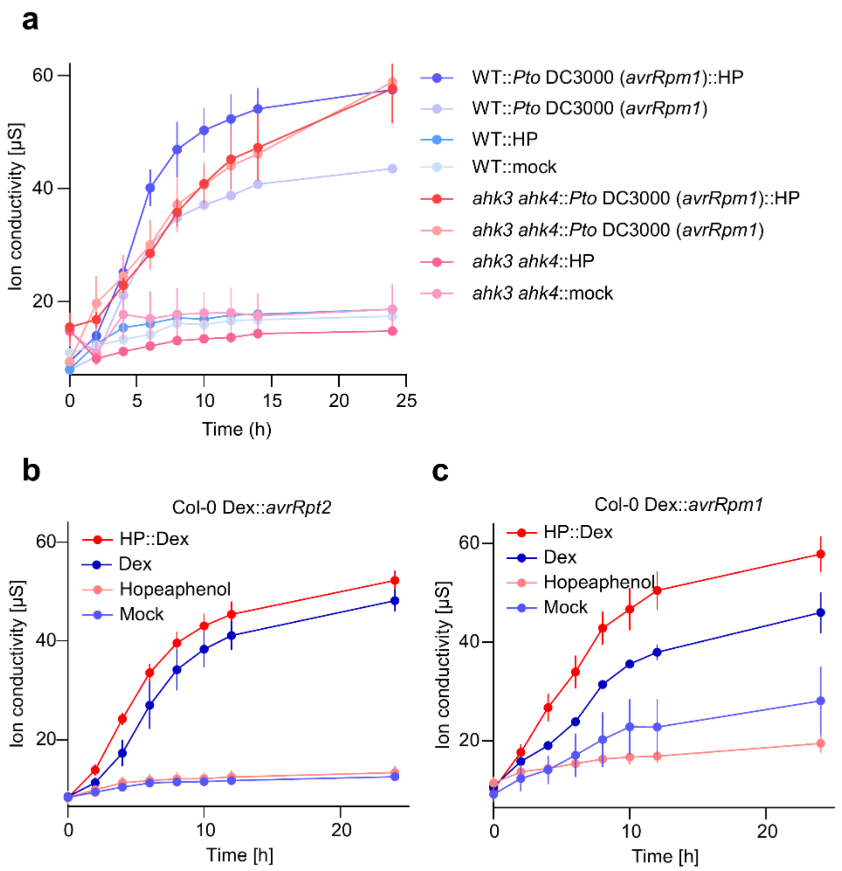


**Extended Data Fig. 6. HP potentiates ETI-associated ion leakage and Dex-inducible effector responses. a,** Ion leakage in WT and *ahk3 ahk4* following mock, HP, *Pto* DC3000 carrying *avrRpm1*, or HP pretreatment followed by *Pto* DC3000 carrying *avrRpm1*. **b,c,** Ion leakage in Col-0 Dex::*avrRpt2* (b) and Col-0 Dex::*avrRpm1* (c) plants following mock, HP, Dex, or HP pretreatment followed by Dex. All ion conductivity was measured every 2 h. Points show mean ± SEM (n = 3).
