## Supplementary Tables for "Hopeaphenol targets histidine kinase signaling across kingdoms to suppress bacterial virulence and potentiate plant immunity"

**Supplementary Table 1.** Arabidopsis lines used in this study.

| Plant line | Allele or transgene | Background | Source or reference |
|---|---|---|---|
| Col-0 | Wild-type | Col-0 | Lab stock |
| *ahk2* | SAIL_1289_G06 | Col-0 | ABRC |
| *ahk3* | GK-105E02 | Col-0 | ABRC |
| *ahk4* | GK-509F06 | Col-0 | ABRC |
| *ahk2 ahk3* | *ahk2-2 ahk3-3* | Col-0 | E. Meyerowitz; Higuchi et al. (2004) |
| *ahk2 ahk4* | *ahk2-2 cre1-12* | Col-0 | E. Meyerowitz; Higuchi et al. (2004) |
| *ahk3 ahk4* | *ahk3-3 cre1-12* | Col-0 | E. Meyerowitz; Higuchi et al. (2004) |
| Dex::*avrRpt2* | Dex::*avrRpt2* | Col-0 | Lab stock |
| Dex::*avrRpm1* | Dex::*avrRpm1* | Col-0 | Lab stock |

**Supplementary Table 2.** Bacterial strains used in this study.

| Strain name | Description | Source |
|---|---|---|
| *Pseudomonas syringae* pv. *tomato* DC3000 | Wild-type strain | Lab stock |
| *Pto* DC3000 ΔCEL | Mutant lacking the conserved effector locus | Lab stock |
| *Pto* DC3000 ΔD36E | Mutant lacking 36 type III effector genes | Lab stock |
| *Pto* DC3000 Δ*hrcC* | Type III secretion-deficient mutant | Lab stock |
| *Pto* DC3000 (*avrRpt2*) | DC3000 expressing AvrRpt2 | Lab stock |
| *Pto* DC3000 (*avrRpm1*) | DC3000 expressing AvrRpm1 | Lab stock |
| *Escherichia coli* BL21 (DE3) | Recombinant protein expression strain | Lab stock |
| *Pectobacterium atrosepticum* BAA672 | Wild-type strain | ATCC (BAA-672) |

**Supplementary Table 3.** Primers used in this study.

| Primer name | Sequence (5’ – 3’) | Description |
|---|---|---|
| F_GW_RhpS | CACCAGCAGCGGCCTGGTGCCGCGCGGCAG CATGATTCGCAGGTTCGACAC | Gateway cloning of *rhpS* (PSPTO_2222) |
| R_GW_RhpS | TCAAAGGCGCGGCAGG | Gateway cloning of *rhpS* (PSPTO_2222) |
| F_RhpS_P40A | GCGCCCCATGGCGCCGCCAAGGC | Q5 site-directed mutagenesis to generate RhpS P40A |
| R_RhpS_P40A | ATGCCGTATTGCGTGAACCAGATGAAGGCCAG CAAG | Q5 site-directed mutagenesis to generate RhpS P40A |
| F_GW_PhoQ | CACCATGATTCGTTCGCTTCGCC | Gateway cloning of *phoQ* (PSPTO_1680) |
| R_GW_PhoQ | TCACTGCGCCAGAAAATGAATC | Gateway cloning of *phoQ* (PSPTO_1680) |
| F_GW_1482 | CACCATGCAAAGCAGAGTCAGGGA | Gateway cloning of PSPTO_1482 |
| R_GW_1482 | CTGGCCAGCGGCAC | Gateway cloning of PSPTO_1482 |
| F_GW_1803 | CACCATGCGCTCATTGTTCTGGC | Gateway cloning of PSPTO_1803 |
| R_GW_1803 | CTAAACCGATGAAGGGCTGA | Gateway cloning of PSPTO_1803 |
| F_GW_CorS | CACCATGAGGCACGACGCTCGCTC | Gateway cloning of *corS (*PSPTO_4705) |
| R_GW_CorS | CTAAAACGGCGCCGGCAC | Gateway cloning of *corS* (PSPTO_4705) |
| F_GW_GacS | CACCATGCTGACCAAGCTGGGTATAAAAGG | Gateway cloning of *gacS* (PSPTO_1691) |
| R_GW_GacS | TCACGCTGTCACCCGC | Gateway cloning of *gacS* (PSPTO_1691) |
| F_GW_ECA4410 | CACCAGCAGCGGCCTGGTGCCGCGCGGCAG CATGATGATACGCGGCCG | Gateway cloning of ECA_RS4410 |
| R_GW_ECA4410 | TTATGCCTTCCAACGAGGTACAC | Gateway cloning of ECA_RS4410 |
| F_ECA4410_P39 A | CAGACACAAAGCTCTCGAGGAGATTGTTG | Q5 site-directed mutagenesis to generate ECA4410 P39A |
| R_ECA4410_P3 9A | TCTCCACCCAGTGAG | Q5 site-directed mutagenesis to generate ECA4410 P39A |
| F_GW_CHASE3 | CACCCATCACAGCAGCGGCCT | Gateway cloning of AHK3 CHASE domain |
| R_GW_CHASE3 | TCACCGCATCTCGTGCTT | Gateway cloning of AHK3 CHASE domain |
| F_CH4_D262A | TGGTGAAGAAGCGAGAGAAAACATCTTAC | Q5 site-directed mutagenesis to generate AHK4 CHASE D262A |
| R_CH4_D262A | GACATCATGTCTAAGCTTTC | Q5 site-directed mutagenesis to generate AHK4 CHASE D262A |
| F_CH4_L284A | GTTCCGGCTCGCGGAGACCCATCAC | Q5 site-directed mutagenesis to generate AHK4 CHASE L284A |
| R_CH4_L284A | GGGGAAGTGAGCACG | Q5 site-directed mutagenesis to generate AHK4 CHASE L284A |
| F_CH4_T278I | GGCCGTGCTCATTTCCCCGTTCC | Q5 site-directed mutagenesis to generate AHK4 CHASE T278I |

| | | |
|---|---|---|
| R_CH4_T278I | TTGCCAGTCTCTCGTGC | Q5 site-directed mutagenesis to generate AHK4 CHASE T278I |

**Supplementary Table 4.** Plasmids used in this study.

| Plasmid name | Description |
|---|---|
| pHisMBP_RhpS | pDEST-HisMBP carrying full-length *rhpS* (PSPTO_2222) for recombinant protein expression |
| pHisMBP_P40A | pDEST-HisMBP carrying full-length *rhpS* P40A for recombinant protein expression |
| pHisMBP_PhoQ | pDEST-HisMBP carrying full-length *phoQ* (PSPTO_1680) for recombinant protein expression |
| pHisMBP_1482 | pDEST-HisMBP carrying full-length PSPTO_1482 for recombinant protein expression |
| pHisMBP_1803 | pDEST-HisMBP carrying full-length PSPTO_1803 for recombinant protein expression |
| pHisMBP_CorS | pDEST-HisMBP carrying full-length *corS* (PSPTO_4705) for recombinant protein expression |
| pHisMBP_GacS | pDEST-HisMBP carrying full-length *gacS* (PSPTO_1691) for recombinant protein expression |
| pHisMBP_4410 | pDEST-HisMBP carrying full-length ECA_RS4410 for recombinant protein expression |
| pHisMBP_4410_P39A | pDEST-HisMBP carrying full-length ECA_RS4410 P39A for recombinant protein expression |
| pHisMBP_CHASE3 | pDEST-HisMBP carrying AHK3 CHASE domain for recombinant protein expression |
| pTwistENTR_CHASE2 | pTwistENTR entry clone carrying the sequence encoding AHK2 CHASE domain; synthesized by Twist Bioscience |
| pTwistENTR_CHASE4 | pTwistENTR entry clone carrying the sequence encoding AHK4 CHASE domain; synthesized by Twist Bioscience |
| pHisMBP_CHASE2 | pDEST-HisMBP carrying AHK2 CHASE-domain sequence transferred from pTwistENTR_AHK2_CHASE for recombinant protein expression |
| pHisMBP_CHASE4 | pDEST-HisMBP carrying AHK4 CHASE-domain sequence transferred from pTwistENTR_AHK4_CHASE for recombinant protein expression |
| pHisMBP_D262A | pDEST-HisMBP carrying AHK4 CHASE domain D262A for recombinant protein expression |
| pHisMBP_L284A | pDEST-HisMBP carrying AHK4 CHASE domain L284A for recombinant protein expression |
| pHisMBP_T278I | pDEST-HisMBP carrying AHK4 CHASE domain T278I for recombinant protein expression |
| pET28a_PKM1 | pET28a carrying full-length PKM1 (Uniprot; P14618) for recombinant protein expression |
